# 15-PGDH Regulates Hematopoietic and Gastrointestinal Fitness During Aging

**DOI:** 10.1101/2020.12.22.424017

**Authors:** Won Jin Ho, Julianne N.P. Smith, Young Soo Park, Matthew Hadiono, Kelsey Christo, Alvin Jogasuria, Yongyou Zhang, Alyssia V. Broncano, Lakshmi Kasturi, Dawn M. Dawson, Stanton L. Gerson, Sanford D. Markowitz, Amar B. Desai

## Abstract

Emerging evidence implicates the eicosanoid molecule prostaglandin E2 (PGE2) in conferring a regenerative phenotype to multiple organ systems following tissue injury. As aging is in part characterized by loss of tissue stem cells’ regenerative capacity, we tested the hypothesis that the prostaglandin-degrading enzyme 15-hydroxyprostaglandin dehydrogenase (15-PGDH) contributes to the diminished organ fitness of aged mice. Here we demonstrate that genetic loss of 15-PGDH (*Hpgd*) confers a protective effect on aging of murine hematopoietic and gastrointestinal (GI) tissues. Aged mice lacking 15-PGDH display increased hematopoietic output as assessed by peripheral blood cell counts, bone marrow and splenic stem cell compartments, and accelerated post-transplantation recovery compared to their WT counterparts. Loss of *Hpgd* expression also resulted in enhanced GI fitness and reduced local inflammation in response to colitis. Together these results suggest that 15-PGDH negatively regulates aged tissue regeneration, and that 15-PGDH inhibition may be a viable therapeutic strategy to ameliorate age-associated loss of organ fitness.

**ARTICLE SUMMARY:** 15-PGDH as a Driver of Age-Related Tissue Dysfunction

## INTRODUCTION

In the process of aging, organs gradually lose their overall fitness. A significant cause of this is decreased regenerative capacity and, thus, the accumulation of damage over time. This loss of capacity results from acquired and genetic factors, and is hastened by both behavioral and environmental factors [1]. One hallmark of aging in mammalian models is stem cell exhaustion, which is characterized by degenerative changes in tissue stem cells, stem cell niches, and signaling pathways responsible for maintaining tissue homeostasis. This process is particularly relevant in organ systems that undergo continuous self-renewal such as the bone marrow (BM) and the gastrointestinal (GI) tract.

Hematopoietic stem cells (HSCs) are derived from the BM and are tasked with ensuring a consistent output of differentiated blood cell types throughout the lifetime of an organism. Self-renewal, differentiation to mature progeny, and proliferation in response to infection, stress, or injury, must be balanced in order to provide lifelong hematopoietic function. At homeostasis, HSCs maintain genomic stability and avoid replicative stress by existing in a quiescent state, enforced by both HSC intrinsic and microenvironmental cues. As an organism ages, however, HSC function declines due to a number of factors, including the accumulation of DNA damage, epigenetic alteration, metabolic changes, loss of cell polarity, exposure to increased basal inflammation, and niche dysfunction [2-8]. As a result, self-renewal capacity declines, myeloid and platelet differentiation predominates at the expense of lymphoid differentiation, and clonal hematopoiesis can emerge [9] [10] [11] [12] [13]. In mouse models, aging is characterized in part by a significantly impaired ability of hematopoietic stem cell populations to engraft and differentiate following transplantation [11] [14] [9], further demonstrating that HSC function declines with age, despite an overall increase in phenotypic HSC number [15] [11].

A potential role for Prostaglandin E2 (PGE2) in modulating hematopoietic function at steady state and post-hematopoietic stem cell transplantation (HST) has been demonstrated by: i) effects of PGE2 on hematopoiesis in zebrafish; ii) increased effectiveness of donor HSCs treated *ex vivo* with PGE2 analogs prior to HST (in both murine models and non-human primates); and iii) protective effects of injected PGE2 in mice receiving sublethal irradiation [16] [17] [18] [19] [20] [21] [22] [23]. Significantly, our work has shown *in vivo* that the prostaglandin-degrading enzyme 15-PGDH plays a major role in regulating BM PGE2 and in regulating a PGE2-controlled pathway of downstream cytokines, including CXCL12 [24] and Stem Cell Factor (SCF) [25], which are key components of the BM stem cell niche [26] [27]. We have characterized highly-potent *in vivo* active 15-PGDH inhibitors, and we have shown that this 15-PGDH inhibition (PGDHi) accelerates hematopoietic regeneration by up to 6 days faster following HST. Notably we have already demonstrated that PGDHi following transplantation of aged donors into aged recipients promotes accelerated hematopoietic recovery, suggesting that activity of this pathway is conserved in the aging hematopoietic system [26].

Over the course of an organism’s lifespan, the GI tract undergoes constant basal regeneration, in part as defense against injurious foreign substances, pathogens, and other insults. Through complete turnover of the intestinal epithelium every 3-5 days, the GI tract is able to maintain proper functioning while sustaining various biochemical and mechanical stress [28]. However, studies utilizing mice models and human samples have suggested that with age, GI fitness declines, as represented by a relative reduction in mucosal heights and accumulation of genetic mutations in its stem cell population. These changes indicate that aging renders the GI tract less functional, with decreased regenerative capacity and increased vulnerability to pathological insults [29-33].

In the GI tract, several studies have clearly demonstrated the importance of PGE2 in maintaining the intestinal stem cell niche, especially in relation to its interaction with the Wnt cascade [34-36]. Furthermore, in the context of inflammatory bowel diseases (IBD), several human epidemiologic studies and animal model experiments have suggested that PGE2 signaling pathways serve to protect and/or repair the GI tract [37-39]. Based on this cumulative evidence, we hypothesized that bolstering PGE2 signaling may augment organ fitness as well as enhance tissue repair in the elderly.

We have previously characterized 15-hydroxyprostaglandin dehydrogenase (15-PGDH), the enzyme responsible for the rate-limiting step of PGE2 degradation, as a potential therapeutic target to augment tissue regeneration in models of bone marrow transplant and IBD [40]; these findings have further established the positive role of PGE2 and the negative role of 15-PGDH in organ self-renewal. Thus, in this study, we explore the hypothesis that 15-PGDH may in fact be a negative regulator of age-related fitness. Indeed, we demonstrate for the first time that genetic ablation of 15-PGDH protects mice from natural age-related changes and improves injury responses in the bone marrow and the colon.

## MATERIALS AND METHODS

### Mice

Mice studies were conducted in the Case Animal Resource Center with animal care and procedures performed under a protocol approved by and in accordance with guidelines of Case Western Reserve University’s Institutional Animal Care and Use Committee. The animals were housed in standard microisolator cages and maintained on a defined irradiated diet (Prolab Isopro RMH 3000) and autoclaved water. 15-PGDH knockout mice were maintained on an FVB background as previously described [41] and were always compared to control FVB wild-type mice. Identities of 15-PGDH knockout and wild-type mice were confirmed by genotyping as previously described [42].

### PGE2 Measurement in Mouse Tissues

Solid tissues were harvested, rinsed in ice-cold PBS containing indomethacin (5.6 μgm/ml), and snap frozen in liquid nitrogen. Marrow was flushed into 1.5ml ice cold PBS with indomethacin, pelleted at 2000 rpm for 5 minutes in an Eppendorf centrifuge, and snap frozen over liquid nitrogen. Frozen colons were pulverized over liquid nitrogen prior to further homogenization. All frozen samples were placed in Eppendorf tubes in cold PBS with indomethacin and were homogenized using a conical tip electric pestle. The suspension was sonicated in an ice water containing bath sonicator for 5 min using cycles of 20 second of sonication with 20 seconds of cooling, followed by centrifugation for 10 minutes at 13,200 rpm. The PGE2 level of the supernatant was measured using a PGE2 ELISA Kit (R&D Systems, cat. # KGE004B) and normalized to total protein concentration measured by BCA assay (Thermo Scientific, cat. #23225) and expressed as ng PGE2/mg total protein. Each sample was assayed in duplicates.

### Peripheral Blood and Marrow Hematopoietic Analysis

Peripheral blood was collected from mice via submandibular bleeding and blood counts were recorded using the Hemavet 950fs (Drew Scientific). Blood counts were tabulated graphically with error bars corresponding to standard error of the means and compared using 2-tailed t-tests. Bone marrow cellularity was determined using hemocytometer under light microscope. Total marrow was incubated with lineage markers (CD3, CD4, B220, Ter119, CD11b) and Sca1+/c-Kit+ cells in lineage negative cells determined via flow cytometry on a BD LSRII instrument (BD Biosciences). Data was analyzed on FlowJo software (Treestar). Values were tabulated graphically with error bars corresponding to standard error of the means and compared using 2-tailed t-tests.

### CFU-S Assay

15 month old 15-PGDH WT and KO were sacrificed and marrow flushed for transplantation into lethally irradiated 8-week old FVB WT mice. 200K total cells were injected via tail vein. At day 12, mice were sacrificed and spleens harvested and weighed. Isolated spleens were fixed in Bouin’s solution for 10 minutes and transferred to 10% formalin for 24 hours. Colony counts were recorded after fixation.

### Hematopoietic Competitive Repopulation Assay

One million whole bone marrow cells from 14 month old WT or 15-PGDH KO mice (CD45.1) were mixed with 6 month old WT FVB (CD45.2) mice at a 1:1 ratio and injected via tail vein into lethally irradiated 2 month old WT FVB CD45.1 recipients. At eight and twenty weeks post-transplant, the mouse chimerism was measured via peripheral blood measuring CD45.1/CD45.2. At twenty weeks, the animals were sacrificed to measure marrow chimerism as well as LSK compartment chimerism using the CD45.1 and CD45.2 markers. Values were tabulated graphically with error bars corresponding to standard error of the means and compared using 2-tailed t-tests.

### Gross Measurements of Small Intestine and Colon Lengths

To measure both small intestine and colon lengths in mice, the distance from the proximal end of the duodenum to the ileocecal junction and the distance from colocecal junction (the point of ileocecal junction) to the rectum were measured, respectively, after rinsing out the luminal debris with ice cold PBS and laying them on a moist non-stick surface.

### Colon Histology and Immunohistochemistry

To perform histological analyses, the harvested colons were fixed in formalin in “swiss-roll” fashion and paraffin embedded. Other organs were also harvested and formalin-fixed paraffin-embedded. Histological morphology of the organs from wild-type and knockout mice were compared upon hematoxylin eosin staining. To characterize the distribution of mucin, PAS/Alcain blue staining was performed. The colons were further analyzed for steady-state proliferative state and degree of damage by immunohistochemical staining of Ki67 and H2AX, respectively. Tissue sections were baked for 40-60 minutes at 60°C followed by deparaffinization and rehydration by brief incubations in xylene and serial ethanol dilutions. Antigen retrieval was performed using Antigen Retrieval Solution (DAKO) at >98°C in steamer. Nonspecific peroxide reaction was blocked by 10 minute in peroxidase block, and nonspecific antibody staining was blocked by Rodent Block M solution (Biocare Medical, Concord, CA). After washing the slides in TBS with Tween (TBS-T), primary antibody incubation with anti-Ki67 rabbit anti-mouse (SP6; ThermoFisher Scientific; 1:200 dilution) was performed for 1 hour at room temperature. For H2AX, sections were incubated with anti-phospho-H2AX rabbit anti-mouse (Ser139; Cell Signaling; 1:450 dilution) overnight at 4°C. Slides were again washed in TBS-T two times followed by incubation with anti-rabbit EnVision detection kit (DAKO) for 1 hour in room temperature. Sections were exposed to DAB and were counterstained in hematoxylin and TBS-T.

### Image Analyses

Images were captured using microscope model via SPOT Ver.5.2 software (Sterling Heights, MI). To measure the crypt heights, ImageJ software was used. Mean crypt height for each mouse was calculated by averaging 30-50 randomly selected full crypt heights for proximal, middle, and distal thirds of the colons (total 90-150 crypts per mouse). Amount of Alcian-Blue stain was quantified by area per mucosal tissue area using thresholding settings on ImageJ. Particle analysis function was used to quantify the number of goblet cells per millimeter length of mucosa. To compare the steady-state proliferative state and degree of DNA damage, the number of Ki67-positive cells and phospho-H2AX-positive cells per crypt were manually quantified in and averaged over approximately 50 random crypts from each mouse. For H2AX quantification, any cell with at least one positive focus was considered positive.

### Colon Injury Model

To examine whether aged KO mice withstand overt colon injury better than the WT, aged-matched male mice were exposed to 2% w/v dextran sodium sulfate (MP Biochemicals) in drinking water for seven days. The matched pairs were co-housed prior to the study. The mice were observed for clinical indices of colonic injury by measuring the weights, stool consistency (scored 0-3), and bleeding severity (scored 0-3) on a daily basis from onset of DSS and for three additional days following the switch to normal drinking water on the eight day of experiment. Mice were sacrificed on day 10 to analyze other markers of colonic injury, i.e. colon lengths, histological scores of crypt destruction, and the levels of inflammatory cytokines.

### Histological Assessment of Cryptitis

To histologically assess crypt destruction along the entire length of the colon, the harvested colons were cut open longitudinally and laid flat inside agar blocks, which were formalin-fixed paraffin-embedded for sectioning. Crypt destruction, i.e. cryptitis, was semi-quantitatively scored by the product of the severity of crypt loss (0: normal, 1: lower one third, 2: lower two thirds, 3: full crypt but intact epithelium, 4: loss of the epithelial layer) and the longitudinal extent of the colon involved (0: 0%, 1: 25%, 2: 50%, 3: 75%, 4: 100%).

### Cytokine Measurements in the DSS-treated Colons

Since DSS colonic injury predominantly affects the distal portion, distal two-centimeter length of each colon was harvested, rinsed in cold PBS, snap frozen, and pulverized over liquid nitrogen. The samples were further homogenized using a conical tip electric pestle in 20mM Tris-buffer containing 150mM NaCl, 1mM EDTA, 1mM EGTA, 1% Triton X-100, 1% phosphatase inhibitor cocktail 2 (Sigma), 1% phosphatase inhibitor 3 (Sigma), and 1X HALT protease inhibitor cocktail (ThermoFisher Scientific). The homogenized samples were centrifuged at 12,000 rpm for 12 minutes, and the resulting supernatants were assayed for specific cytokines using the mouse pro-inflammatory cytokine panel detected via Mesoscale SECTOR® Imager 2400 multiplex analyzer (Mesoscale Discovery) following the manufacturer’s protocol. Levels of cytokines were normalized to the total amount of protein in the samples quantified by BCA assay (pg cytokine/mg total protein).

## RESULTS

### Aged 15-PGDH KO mice display elevated PGE2 in multiple organs

We have previously reported that basal PGE2 doubles in young (2-3 month) mice lacking 15-PGDH [27]. To determine if the loss of 15-PGDH also results in elevated PGE2 in aged mice, PGE2 levels were measured in multiple organs of aged (15 month) wildtype littermate (WT) and 15-PGDH knockout (KO) mice. PGE2 levels in aged WT mice were consistent with our previously published analysis of young WT animals from the same colony [27]. Aged 15-PGDH KO mice demonstrated a similar increase in PGE2 levels as observed in young KO versus WT animals, with bone marrow (1.77 fold), spleen (1.65 fold), lung (1.39 fold), and colon (1.83 fold) all demonstrating significant increases in total PGE2 when compared to aged-matched, gender-matched WT counterparts (**Figure 1A-D**). This indicates that genetic loss of 15-PGDH results in a consistent increase in PGE2 levels throughout the lifetime of the animal.

**Figure 1.**
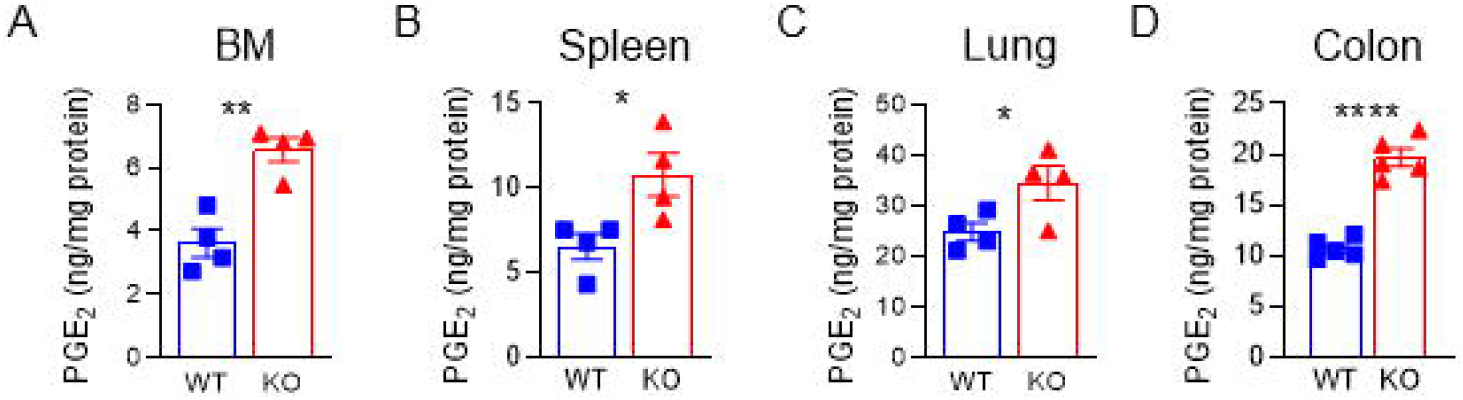
Aged PGDH KO mice display elevated PGE2 (and reduced inflammatory cytokines). (**A-D**) PGE2 levels were quantified in bone marrow (BM), spleen, lung, and colon tissue lysates obtained from 12-15 month old FVB wildtype littermates (WT), and 15-PGDH knockout (KO) mice. N = 4 mice/group.

### White blood cells and HSPCs are increased in aged 15-PGDH KO mice

Young 15-PGDH KO mice display significantly increased total neutrophil counts over WT littermate controls [27]. Here, we examined peripheral blood counts in an aged KO and WT colony (19-23 month). As expected, aged mice displayed higher absolute neutrophil counts compared to young mice from the same colony [27], and consistent with our findings in younger mice, white blood cells (WBCs) and neutrophils (NE) were significantly increased in aged mice lacking 15-PGDH (**Figure 2A-B**). Aging was associated with a decrease in lymphocyte counts when aged WT mice were compared to young animals in the same colony [27], however, aged KO mice demonstrated a notable trend toward increased lymphocytes (P = 0.06) when compared to aged WT controls (**Figure 2C**). We also examined a colony of middle-aged (13-15 month) KO and WT mice and also observed significantly increased neutrophils counts (data not shown). Analysis of other blood cell compartments revealed no impact of 15-PGDH loss on platelets and a decrease in red blood cells and hemoglobin (from 13.4g/dL to 12.2g/dL; **Supplemental Figure 1A**). These results establish that in the absence of 15-PGDH, greater numbers of mature neutrophils and immune cells are maintained in the peripheral blood, even in extremely aged mice.

**Figure 2.**
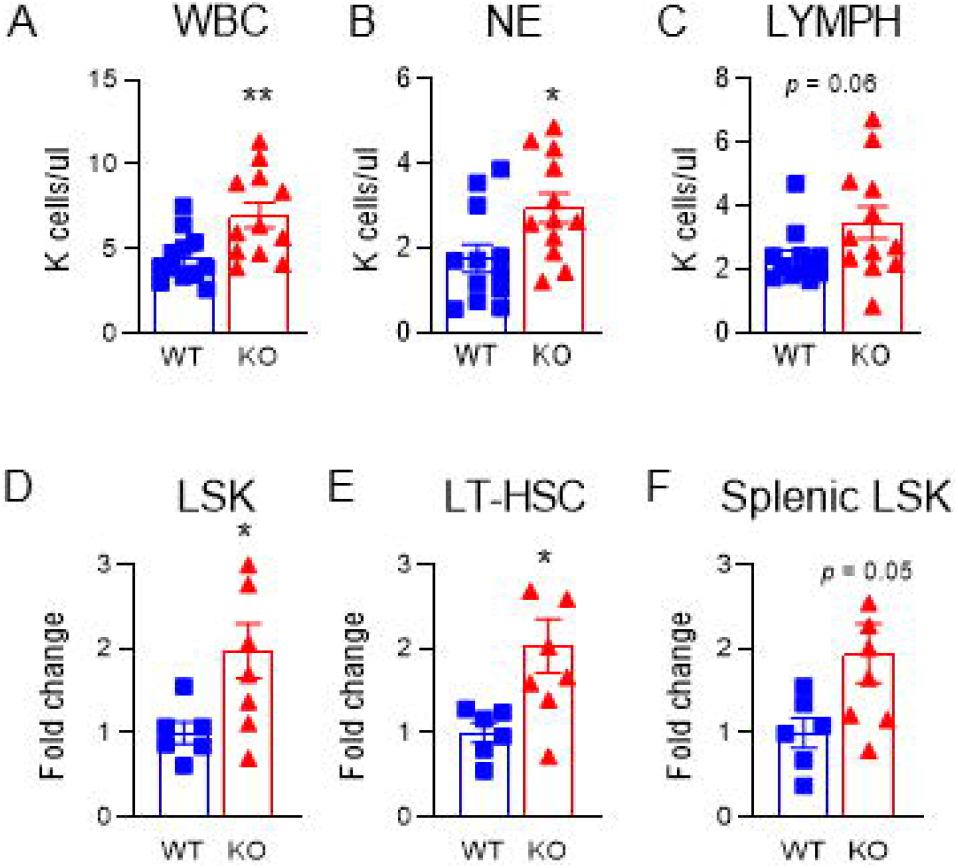
White blood cells and HSPCs are increased in aged PGDH KO mice. (**A-C**) White blood cell (WBC), neutrophil (NE), and lymphocyte (LYMPH) levels were measured in the peripheral blood of aged (19-23 months old) WT and PGDH KO mice by complete blood count analysis. N = 12 mice/group. (**D-F**) Fold change in the number of BM Lin^-^ c-Kit^+^ Sca1^+^ (LSK cells), and Lin^-^ c-Kit^+^ Sca1^+^ CD48^-^ CD150^+^ (phenotypic LT-HSCs) in aged (14-20 months old; median age: 18mos.) PGDH KO relative to WT littermates. (**H**) Fold change in the number of splenic LSK cells in aged (14-20 months old; median age: 18mos.) PGDH KO relative to WT littermates. N = 6-8 mice/group, 2 independent experiments.

To determine if aged 15-PGDH KO mice display any changes in hematopoietic tissues that may underlie the increase in circulating WBCs, we next analyzed the BM and spleen of aged animals. Although the BM cellularity of aged mice lacking 15-PGDH was not significantly different (**Supplemental Figure 1B**), loss of 15-PGDH altered the marrow composition with significant increases observed in the numbers of Ly6C^hi^ monocytes, neutrophils, and CD4+ T lymphocytes, as well as trends towards increased resident macrophages, and CD8+ T lymphocytes (**Supplemental Figure 2**).

To directly determine whether hematopoiesis is impacted by the loss of 15-PGDH, we quantified HSCs in the marrow and spleen of aged KO mice. In spite of the unchanged BM and splenic cellularity, a significant 2-fold increase was observed in both the hematopoietic stem and progenitor cell (HSPC) pool as a whole, identified as Lineage^-^ Sca1^+^ c-Kit^+^ (LSK) cells, and in a subset enriched for long-term HSCs (LT-HSCs; identified as LSK CD48^-^ CD150^+^ cells), was observed in the marrow (**Figure 2D-E**). Moreover, the spleens of aged KO mice also displayed a 1.9-fold increase in hematopoietic stem and progenitor cells (**Figure 2F;** *P* = 0.05). Colony-forming analysis showed trends towards increased frequencies of myeloid and erythroid progenitors in the marrow and spleen of aged 15-PGDH KO mice over WT littermate controls (**Supplemental Figure 3**). Collectively, these data demonstrate that loss of 15-PGDH results in expansion of the phenotypic HSPC pool, which correlates with an increase in circulating white blood cells.

### Loss of 15-PGDH increases CFU-S post-BMT

To more rigorously compare the functional stem cell capacity of bone marrow from 15-PGDH WT and KO aged mice, we quantitated their respective abilities to generate hematopoietic colonies in vivo when used as donors in BM transplantation assays. In this assay, hematopoietic colonies arising in the spleens of transplant recipients were scored 12 days post-transplant. In a BM transplant, hematopoietic stem and progenitor cells must first home and engraft in the recipient spleen where they proliferate to form macroscopic colonies containing myeloid, erythrocytic, granulocytic, and megakaryocytic lineages, all deriving from a single hematopoietic stem cell. To determine whether genetic ablation of 15-PGDH impacts the capacity of aged hematopoietic stem and progenitor cells to appropriately home and engraft, we analyzed colony forming units spleen (CFU-S) post-transplant. Although aged mice did not demonstrate a reduction in CFU-S compared to young counterparts [26], we found that the BM of aged (15 month) PGDH KO mice demonstrates a significant 1.38-fold increase in the total number of CFU-S 12 days post-transplant (**Figure 3A-B**) and a 1.25-fold increase in total spleen weight (**Figure 3C**), indicating that the homing and early engraftment capacity of aged KO HSPC is superior to that of WT. Thus, while a standard feature of hematologic aging is the progressive impairment of aged HSC repopulating capacity (reviewed in [43]), these data suggest that age-related HSC dysfunction may be reversed in mice deficient in 15-PGDH.

**Figure 3.**
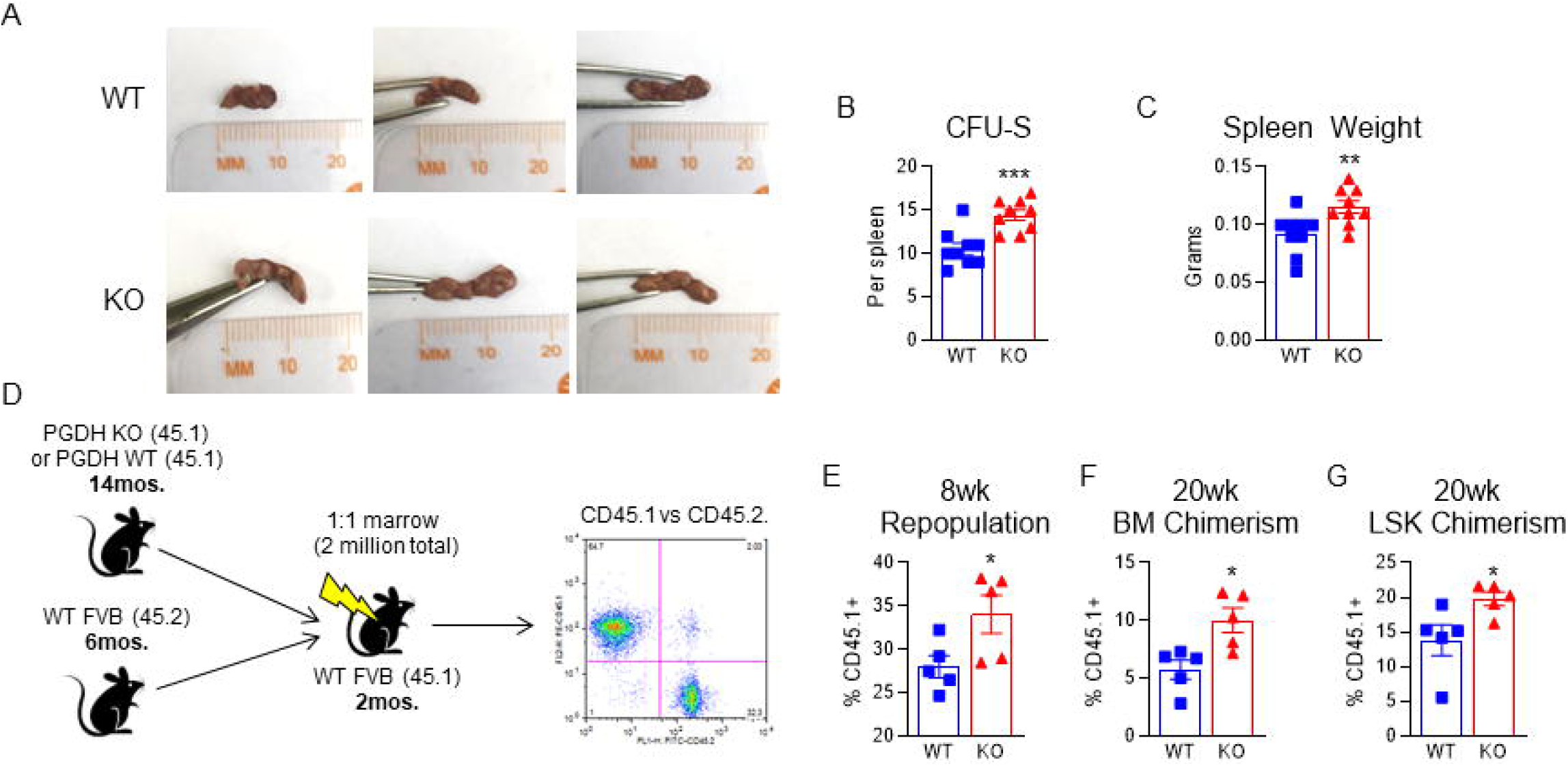
Aged 15-PGDH KO marrow shows enhanced engraftment and hematopoietic repopulation. (**A**) Representative gross images of the spleen of WT recipient mice 12 days post-transplantation of aged (15 months old) WT or PGDH KO BM. (**B**) Hematopoietic colony forming units on the spleen (CFU-S) were scored. N = 9 mice/group. (**C**) Day 12 recipient spleen weights were measured. (**D**) Schematic depicts the competitive transplantation of aged (14 months old) CD45.1+ PGDH KO or WT littermate control BM with the BM of young (6 months old) CD45.2+ competitor mice. (**E**) Peripheral blood repopulation 8wks post-transplant was analyzed as percent CD45.1+ chimerism in peripheral blood in mice that received either WT control or PGDH KO BM. (**F**) BM chimerism was analyzed 20wks post-transplant as the percent CD45.1+ of total BM cells. (**G**) LSK chimerism was analyzed 20wks post-transplant as the percent CD45.1+ of BM LSK cells. N = 5 mice/group.

### The BM of aged 15-PGDH KO mice displays increased repopulating activity

Based upon the observed increase in CFU-S capacity, we next assessed the long-term repopulating capacity of the BM of aged (13 month) PGDH KO mice by competitive transplantation (**Figure 3D**). Although the BM from 6-month-old competitor mice outcompeted donor marrow from either aged WT or KO mice, analysis of peripheral blood chimerism 8 weeks post-transplant revealed a significant advantage of BM from KO animals (34.1±2.2% vs. 28.0±1.3%; **Figure 3E**). When the mice were sacrificed at week 20, peripheral blood cells derived from aged BM comprised only 20% of the recipient peripheral blood and levels were not significantly impacted by 15-PGDH (data not shown). However, aged KO cells demonstrated superior total BM reconstitution (1.75 fold greater representation; **Figure 3F**) and, strikingly, also showed significantly superior reconstitution of the marrow LSK compartment (1.43-fold greater representation, **Figure 3G**). These data suggest that loss of 15-PGDH confers an intrinsic survival and proliferation advantage to aged HSPCs, which results in enhanced fitness of the stem cell pool upon transplantation.

### Loss of 15-PGDH does not result in the accumulation of DNA damage in the BM

DNA damage and impaired DNA damage repair pathways underlie aspects of HSC aging (reviewed in [44]). Additionally, rounds of HSC proliferation can result in replication stress and accumulation of DNA damage [2]. To determine whether increased hematopoietic output observed in aged 15-PGDH KO mice leads to DNA damage, we analyzed phosophorylation of histone H2AX by immunohistochemistry. We found no evidence of DNA damage accumulation (**Supplemental Figure 4**), indicating that the enhanced activity of HSPCs in aged 15-PGDH KO mice is not correlated with DNA damage.

### Mice lacking 15-PGDH exhibit superior colonic fitness with age

Colon crypt height is a measure of colonic fitness as the intestinal epithelium is constantly undergoing repair [31], and is closely linked with intact absorptive function. Therefore, we compared colonic crypt height in aged WT and KO mice. Consistent with our hypothesis, in mice between 10-23 months of age, crypt height was significantly higher in KO versus WT mice (P = 0.003; **Figure 4A-C**). However, Ki67 expression was not significantly different, suggesting that increased crypt height is not due to enhanced steady-state proliferation as a result of lacking 15-PGDH (**Supplemental Figure 5**). This is consistent with the known pro-proliferative impact of PGE2 specifically in the context of injury [40]. Furthermore, DNA damage as assessed by phospho-H2AX foci showed was not significantly different between WT and KO mice (**Supplemental Figure 6**). Since PGE2 is known to upregulate intestinal mucin expression and exocytosis, and mucin plays an important protective role against epithelial injury [45-47], mucin content was also measured in the colons of aged mice. While 15-PGDH did not impact mucin levels in the proximal colon (**Supplemental Figure 7**), KO mice demonstrated approximately two-fold higher levels of PAS/Alcian Blue staining in the distal colon (**Figure 4D-F**), suggesting 15-PGDH KO mice are protected from age-associated loss of distal colon barrier function. Based on our previous observations that genetic knockout as well as chemical inhibition of 15-PGDH provides resistance against colonic injury in mice [40], the current findings indicate that the lack of 15-PGDH benefits overall colon fitness upon aging. This increase in colonic fitness likely owes to enhanced protection from and self-renewal in response to tissue damage over time, rather than increased steady-state proliferation.

**Figure 4.**
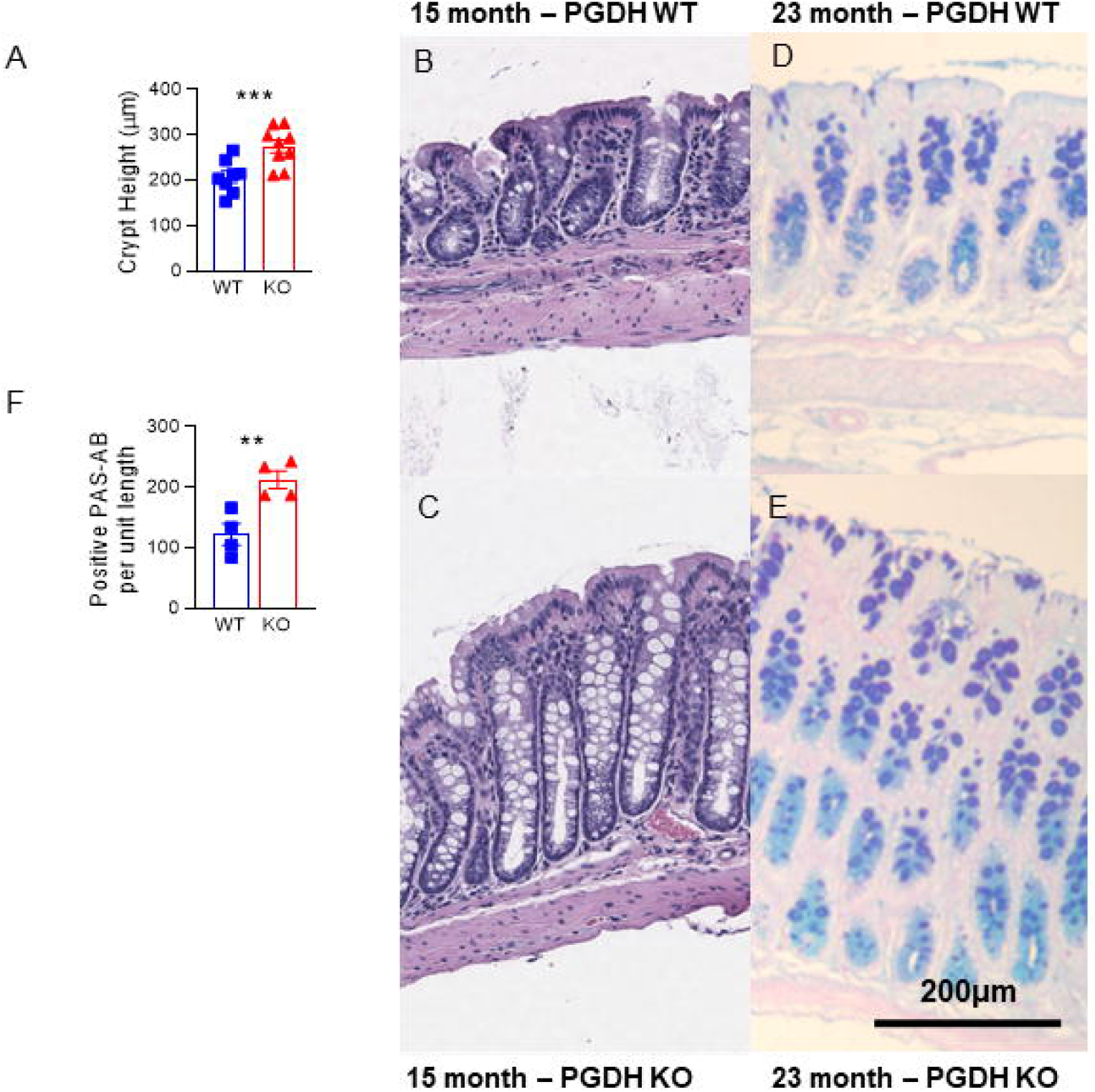
Crypt fitness is superior in aged PGDH KO mice. (**A**) Bar graphs represent crypt heights compared between WT and PGDH KO mice (10-23 months of age; N = 8-9 mice/group). P=0.003 for unpaired t-test. (**B-C**) Representative histologic images of murine colon crypts. H&E for WT and PGDH KO are shown. Scale bar: 200µm. (**D-E**) Representative PAS-Alcian Blue stains performed to visualize mucin content in WT and PGDH KO distal colons are shown. (**F**) Positive PAB-Alcian Blue stains are quantified per unit length for mice with at least 12 months of age. N = 4 mice/group. P=0.008 for unpaired t-test.

### Aged 15-PGDH KO mice demonstrate superior resistance against overt colonic injury

As an alternative test of tissue fitness, we compared response to dextran sodium sulfate (DSS)-induced colon injury in aged 15-PGDH WT versus KO animals. In previous studies, we had demonstrated that 10-12 week old 15-PGDH KO mice were highly resistant to mucosal injury in this DSS-induced colitis model, showing less tissue damage and faster tissue recovery [27]. To determine if this increased mucosal fitness was sustained in aged mice, we administered DSS to 17-23 month-old male mice using a schedule of 7 days administration of DSS, followed by three days of recovery prior to sacrifice (**Figure 5A**). Aged WT mice suffered persistent weight loss (**Figure 5B**) and elevated clinical indices of disease activity, as scored by weight loss, stool consistency, and rectal bleeding (**Figure 5C**), and both weight loss and disease activity were strongly ameliorated in KO mice. Moreover, reduction of colon length is a key index of integrated colitis activity in the DSS model, and loss of 15-PGDH was associated with markedly better retention of colon length on study day 10, with a 1.9 centimeter increase in this measure in KO relative to WT (P=0.01; **Figure 5D;** and **Supplemental Figure 8**). Histologically, cryptitis in the distal colons was also ameliorated in KO mice (P=0.001; **Figure 5E;** and **Supplemental Figure 9**). These findings correlated with reduced inflammation in the colons of KO mice, as measured by reduced TNFα and IL1β (**Figure 5F-G**), as well as trends towards reductions in CXCL1, IL6, and IFNγ, (**Supplemental Figure 10**). As to the enhanced protection in the aged KO mice, the higher overall mucin content observed in the distal colons of aged KO mice provides one potential underlying mechanism. Given that DSS colitis induces injury principally in the distal colon, we also evaluated the number of goblet cells in the distal colon (cells positive for PAS/Alcian Blue) as yet another metric for the differences between the aged KO versus WT mice in their baseline ability to respond to injury. Consistent with the superior resistance of KO mice to DSS colitis, KO mice exhibited nearly two-fold higher numbers of goblet cells per length of mucosa in the distal colon (**Figure 5H**). In summary, by every measure of disease activity in the DSS colitis model, elderly (17-23-month-old) 15-PGDH KO mice maintained the resistance to colon injury that was initially demonstrated by young KO mice.

**Figure 5.**
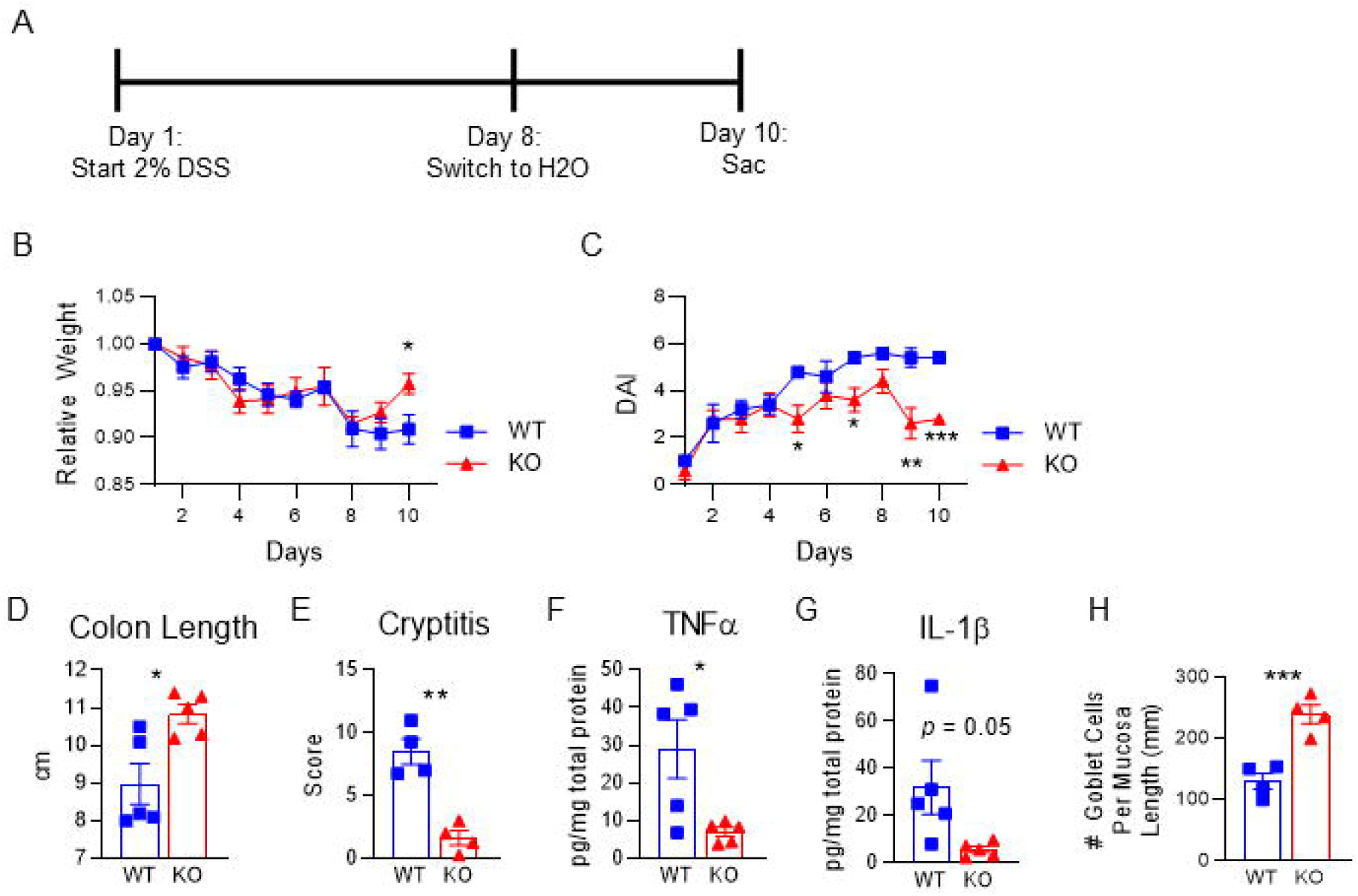
Aged PGDH KO mice show less severe DSS colitis. **(A)** Experimental schema for assaying DSS colitis. **(B)** Mouse weights relative to day one of the experiment across time are shown for WT and PGDH KO mice. Mean±SEM, N = 5 mice/group, *P<0.05, unpaired t-test. **(C)** Disease activity indices (DAI) for the DSS colitis are shown for WT and PGDH KO mice. Mean±SEM, N = 5 mice/group, *P<0.05, unpaired t-test. **(D)** Gross colon lengths resulting from DSS exposure on day 10 are shown, Mean±SEM, N = 5 mice/group, *P=0.01, unpaired t-test. **(E)** Cryptitis scores from histological analysis are shown. Mean±SEM, N = 4 mice/group. ***P=0.001, unpaired t-test. **(F)** TNFα and (G) IL1β inflammatory cytokine content within the distal colons (pg/mg total protein) are quantified. P-values by unpaired t-tests. Mean±SEM, N = 5 mice/group. **(H)** Number of goblet cells per mucosa length (mm) in the distal colon is quantified. ***P=0.001, unpaired t-test. Mean±SEM, N = 4 mice/group.

## DISCUSSION

In summary, we have shown that loss of 15-PGDH enhances hematopoietic and GI fitness with age as demonstrated through enhanced steady-state hematopoiesis, an increase in hematopoietic stem cell engraftment and proliferation post-transplant, and a more robust healing of mucosal injury in the colon. Along with our recent discovery that inhibition of 15-PGDH results in a more robust tissue stem cell proliferation in models of injury [27], 15-PGDH may in fact function as a negative regulator of age-related stem cell fitness. Thus, inhibiting 15-PGDH and, in turn, increasing PGE2 becomes a potential approach not only to better maintain age-related organ fitness but also to treat disease conditions affected by age. Indeed, during the preparation of this manuscript, Blau and colleagues published that 15-PGDH inhibition (PGDHi) increases muscle mass and exercise performance in murine models of sarcopenia. This independent validation of PGDHi efficacy in aging models is an exciting development demonstrating the wide applicability of this strategy [48].

Our data demonstrate that aged 15-PGDH KO bone marrow shows both a short-term increase in hematopoietic engraftment and proliferation, via the CFU-S assay, as well as a long-term competitive advantage in engraftment, via the 20-week competitive repopulation assay. These datasets strongly imply that the number of functional HSCs in the 15-PGDH KO mouse have higher fitness and engraftment potential than their WT counterparts. Aged HSCs also demonstrate a shift in differentiation capacity towards the myeloid lineage; with decreased output of lymphoid cells, aging is associated with overall functional decline of the adaptive immune system [10] [49] [13]. Our results show that 15-PGDH KO mice demonstrate a sustained increase in myeloid with no decrease in lymphoid cell types with age compared to the WT mice, suggesting that the KO HSC population better maintains its differentiation potential to produce both lineage cell types. Although our data do not directly address whether 15-PGDH negatively regulates hematopoiesis via HSC-autonomous or microenvironmental actions, BM lacking 15-PGDH maintains a functional advantage upon transplantation, and thus 15-PGDH-dependent regulation does not require constant involvement of non-hematopoietic cell types. Human BM transplant recipients are often aged individuals receiving either autologous transplants or allogeneic transplants from an aged sibling donor, and increasing age is a significant risk factor in treatment-related mortality and acute and chronic graft-versus-host disease [50] and in poor long term prognosis. These studies demonstrate that marrow of aged 15-PGDH KO mice is more robust in supporting BM engraftment and suggest that the small molecule 15-PGDH inhibitor, SW033291, that we have previously reported, may be of value in supporting BM transplantation when either the donor or recipient is elderly.

Moreover, several conditions associated with hematopoietic aging, including myeloproliferative disorders and myelofibrosis, are typified by the emergence of clonal proliferations with a growth advantage over normal HSCs. It will be intriguing in future studies to examine whether strategies to inhibit 15-PGDH would in these disease contexts restore a growth advantage to normal hematopoietic stem and progenitor cells and their progeny.

In the GI tract, aging is associated with declining self-renewal capacity [31, 32, 51-53]. Consistent with the protective and proliferative effects of PGE2 on the GI epithelium associated with both mucin production and Wnt signaling [34], we find that aged 15-PGDH knockout mice markedly resisted against and recovered faster from colitic injury than did aged wild-type mice. While we have previously made similar observations in young mice, these new observations support that 15-PGDH remains a significant negative regulator of tissue regeneration after injury in aged mice. In humans, the incidence of IBD in the elderly has been steadily increasing, with approximately 15-20% of IBD cases now in individuals over 65 years of age [54]. Elderly IBD patients are at higher risk for hospitalization, longer stays, and worse outcome [54]. Moreover, elderly IBD patients show greater adverse effects from systemic steroids and less benefit from standard therapies [54]. The present studies support that small molecule 15-PGDH inhibitors, which are currently in preclinical development, may offer an alternative strategy for treated IBD in the therapeutically challenging setting of the elderly patient.

In summary, these findings support that 15-PGDH continues to play a significant role as a negative regulator of tissue regeneration throughout life, from both young age through elderly age in mouse models, and moreover, that inhibiting 15-PGDH provides a therapeutic target for reversing some aspects of tissue decline and capacity for repair in the aging organism. The preclinical development of small molecule 15-PGDH inhibitors provides the potential that ultimately this model may be testable in humans.

## Acknowledgments

This work was supported by NIH grants R35 CA197442, R00 HL135740, and T32 EB005583, and by the Radiation Resources Core Facility (P30CA043703), the Hematopoietic Biorepository and Cellular Therapy Core Facility (P30CA043703), the Tissue Resources Core Facility (P30CA043703), and the Cytometry & Imaging Microscopy Core Facility of the Case Comprehensive Cancer Center (P30CA043703).

## Supplementary Figure Legends

**Supplementary Figure 1.**
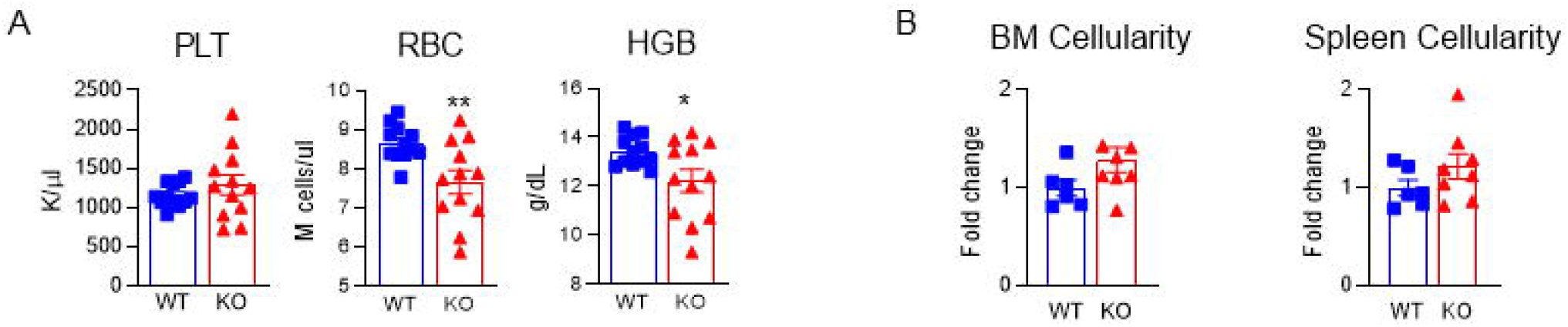
Loss of 15-PGDH does not impact aged marrow and spleen cellularity. **(A)** Platelet (PLT), red blood cell (RBC), and hemoglobin (HGB) quantification by complete blood count analysis in aged (19-23 months old) WT and PGDH KO mice. N = 12 mice/group. **(B)** Fold change in the BM cellularity per hindlimb and cellularity per spleen of aged (14-20 months old; median age: 18mos.) PGDH KO mice relative to WT littermates. N=6-8 mice/group.

**Supplementary Figure 2.**
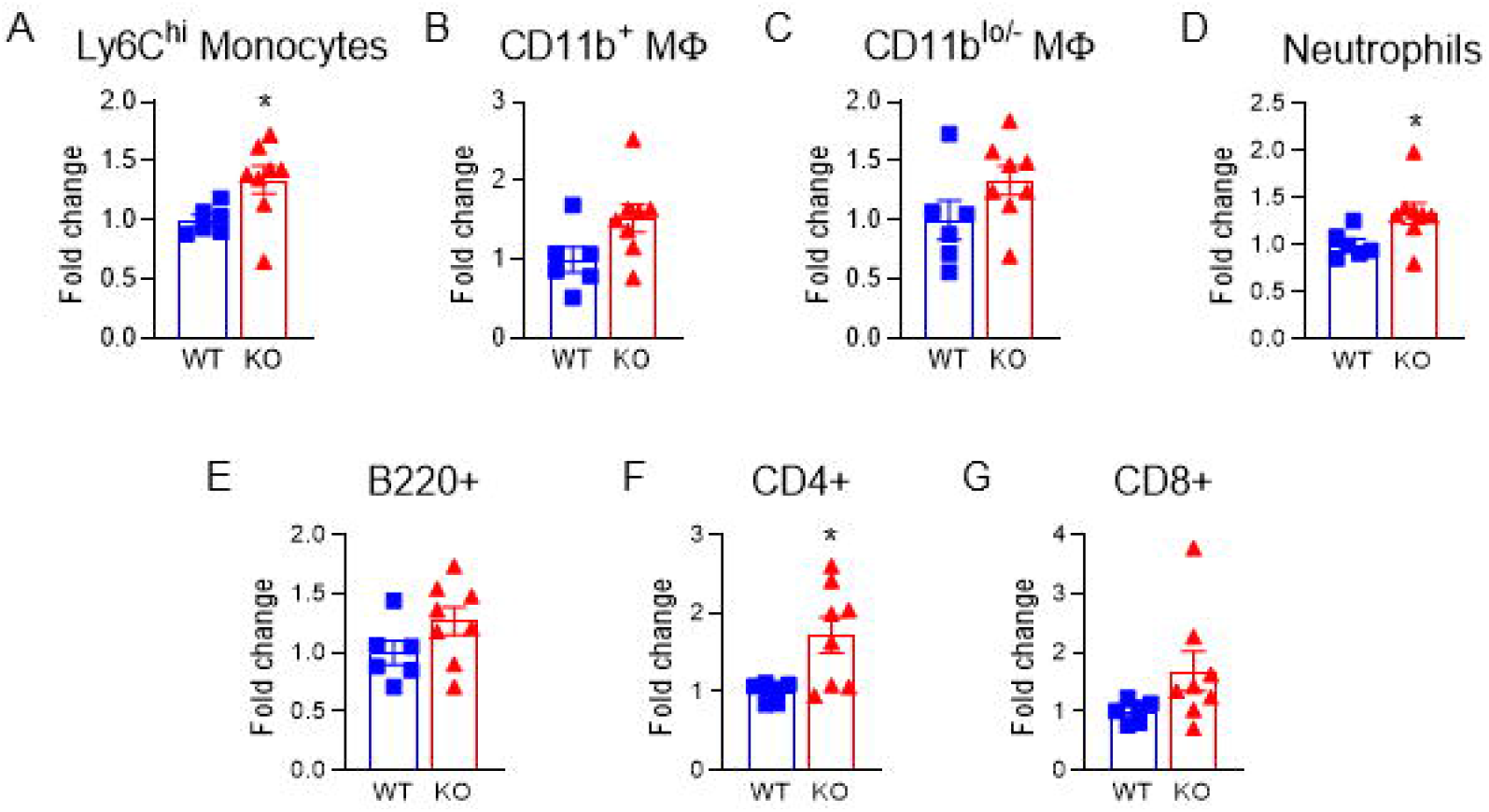
Monocytes, neutrophils, and CD4 T lymphocytes are increased in the marrow of aged PGDH KO mice. (**A-D**) Fold change in the numbers of myeloid cell populations in the BM of aged (14-20 months old; median age: 18mos.) PGDH KO relative to WT mice: Ly6C^hi^ monocytes (defined as CD11b^+^ Ly6C^hi^ cells), CD11b^+^ macrophages (MΦ; defined as CD11b^+^, F4/80^+^, Ly6C^lo^, SSC^lo^ cells), CD11b^lo/-^ MFs (defined as CD11b^lo/-^, F4/80^+^ SSC^lo^ cells), and neutrophils (defined as CD11b^+^ F4/80^-^ Ly6G^hi^ cells). (**E-G**) Fold change in the numbers of lymphoid cell populations in the BM of aged PGDH KO relative to WT mice: B220+ cells (identified as CD11b^-^ B220^+^ cells), CD4+ T lymphocytes (identified as CD11b^-^, CD3e^+^, CD4^+^ cells), and CD8+ T lymphocytes (identified as CD11b^-^, CD3e^+^, CD8^+^ cells). N = 6-8 mice per group.

**Supplementary Figure 3.**
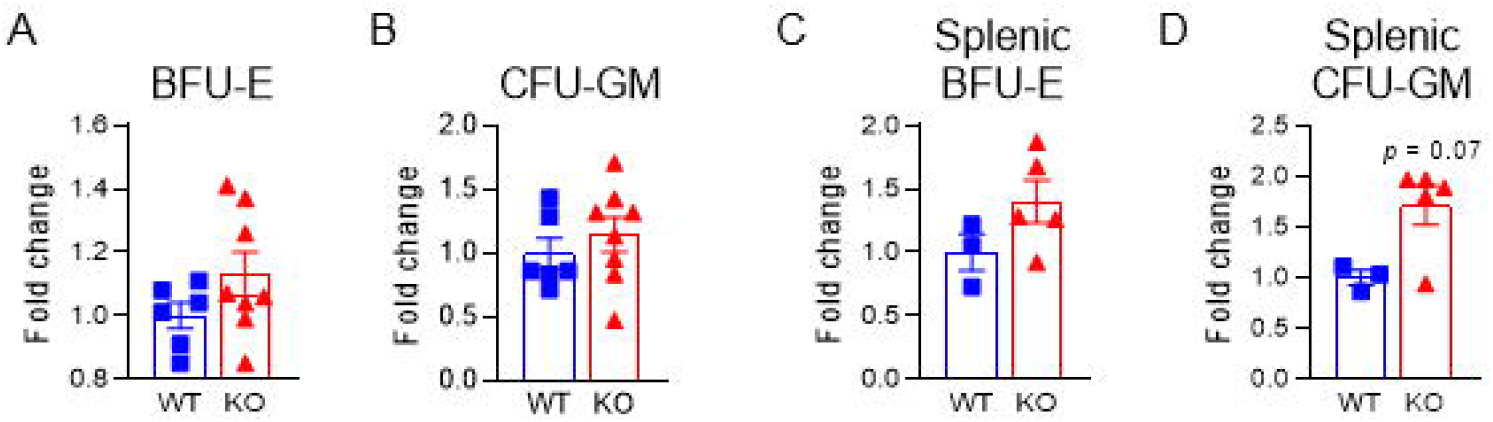
Aged 15-PGDH KO marrow and spleen displays trends towards increased myeloid and erythroid colony-forming activity. (**A-B**) Fold change in the numbers of burst-forming units-erythroid (BFU-E) and colony-forming units granulocyte-macrophage (CFU-GM) derived from the BM of aged (14-20 months old; median age: 18mos.) PGDH KO relative to WT littermate control mice. (**C-D**) Fold change in the numbers of BFU-E and CFU-GM derived from the spleen of aged (14-20 months old; median age: 18mos.) PGDH KO relative to WT littermate control mice. N = 3-8 mice/group.

**Supplementary Figure 4.**
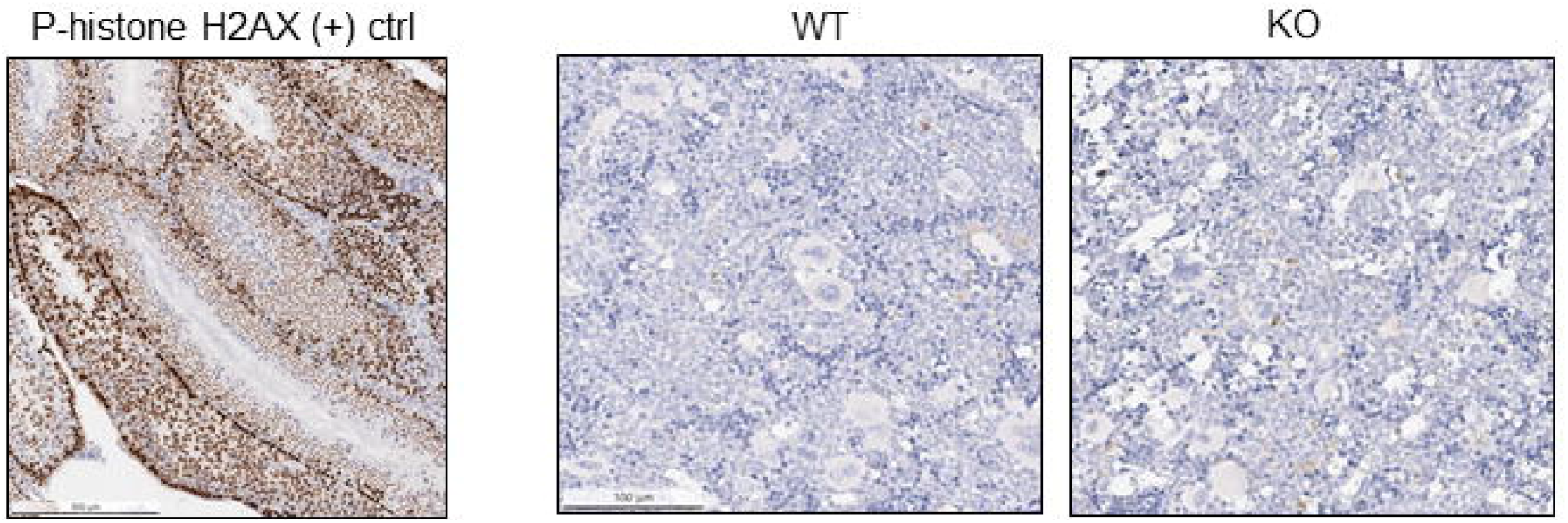
Hematopoietic DNA damage does not accumulate in steady-state BM of aged PGDH KO or WT littermate control mice. Representative images of immunohistochemical phosphorylated histone H2AX staining (brown) in the murine testes as a positive control (left) and in hindlimb BM sections from aged (21-22 months old) WT and PGDH KO mice (right).

**Supplementary Figure 5.**
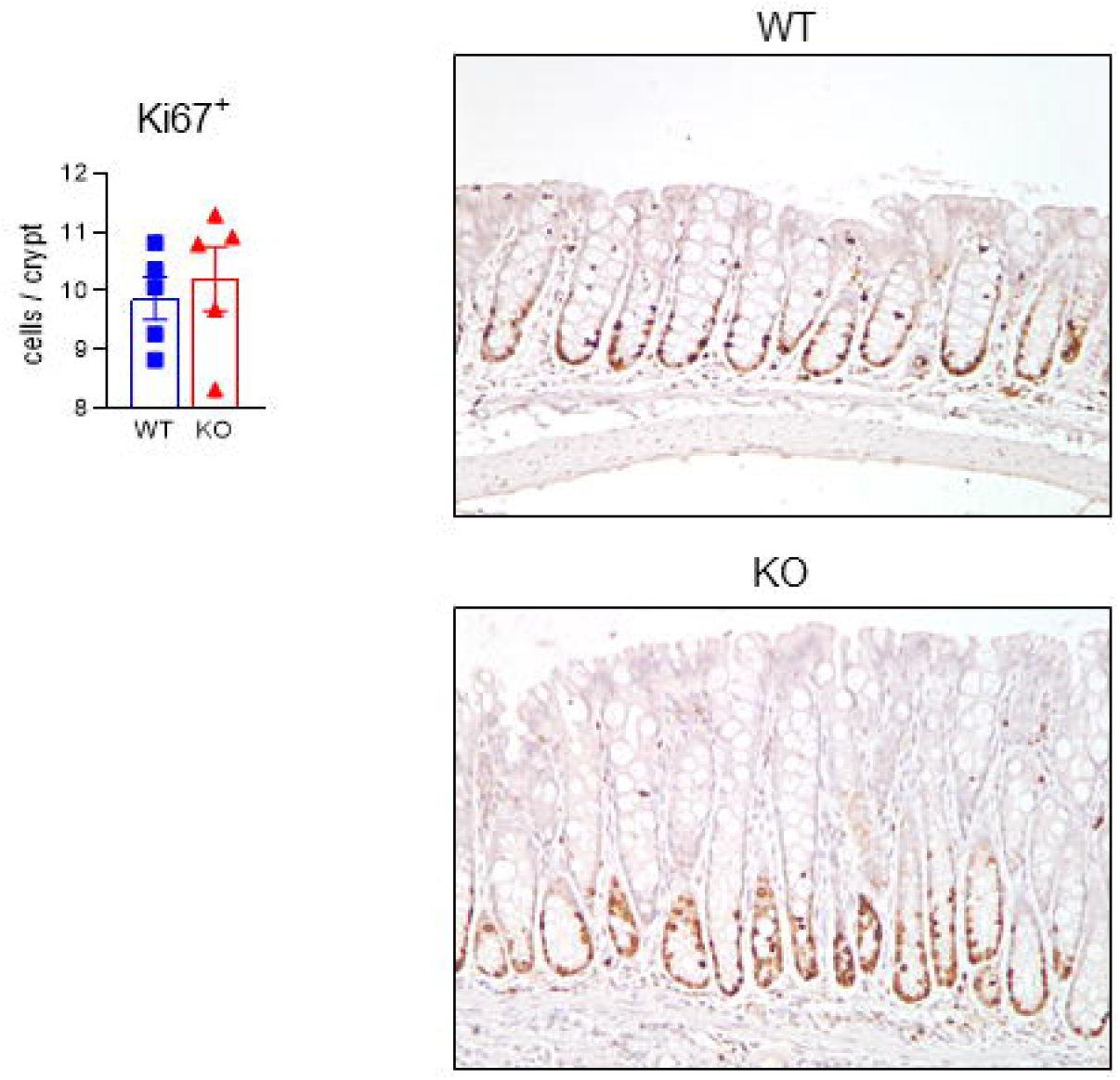
Loss of PGDH does not impact steady-state epithelial cell proliferation. Quantification of Ki67 by immunohistochemistry for aged (≥12 months of age) mouse colons are shown. Mean±SEM, N = 5 mice/group (50-60 crypts per mouse quantified). Representative staining results for both groups are shown.

**Supplementary Figure 6.**
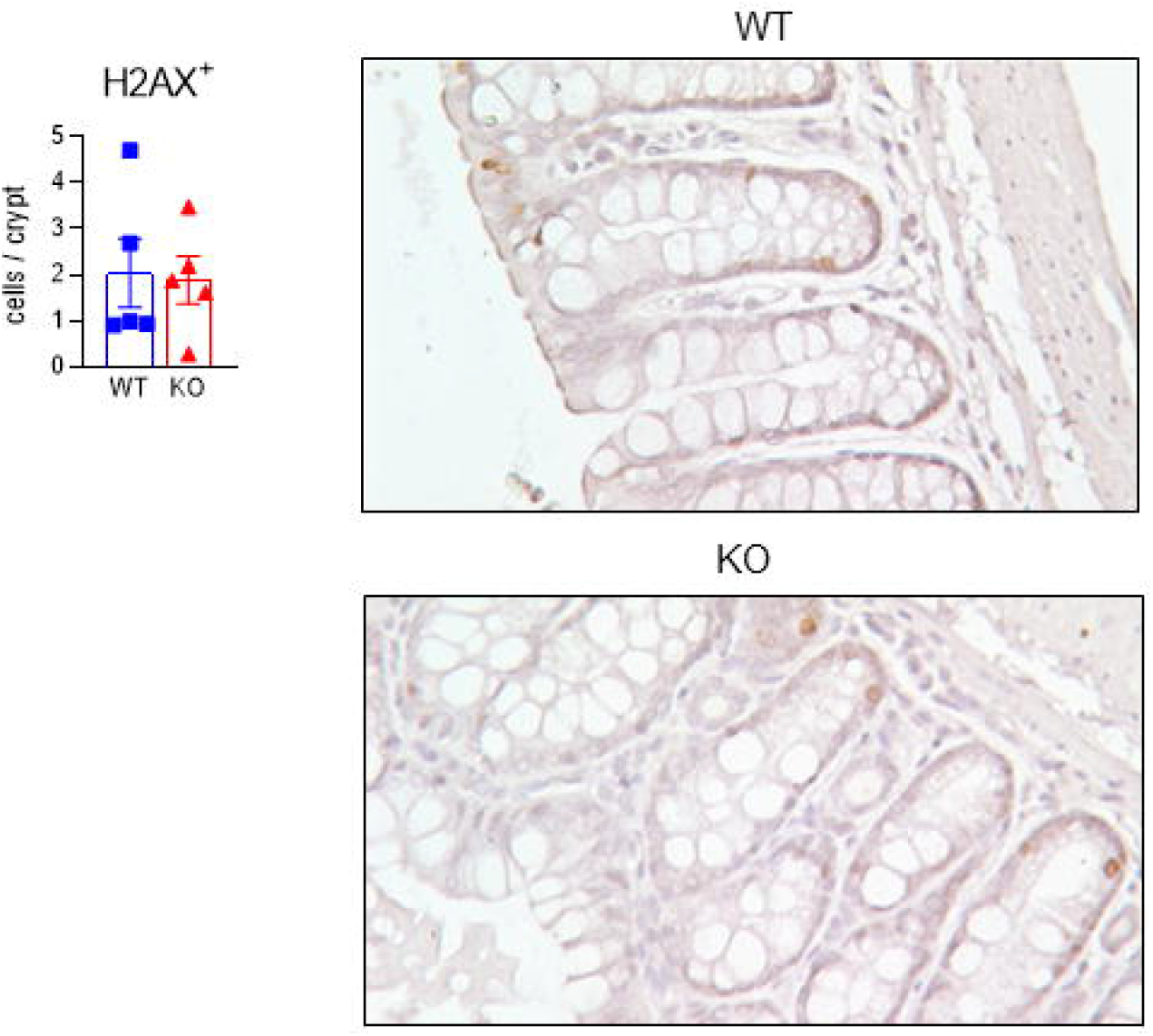
PGDH does not influence the accumulation of DNA damage in steady-state colonic crypts. Quantification of *γ*H2AX by immunohistochemistry for aged (≥12 months of age) mouse colons are shown. Mean±SEM, N = 5 mice/group (50-60 crypts per mouse quantified). Representative staining results for both groups are shown.

**Supplementary Figure 7.**
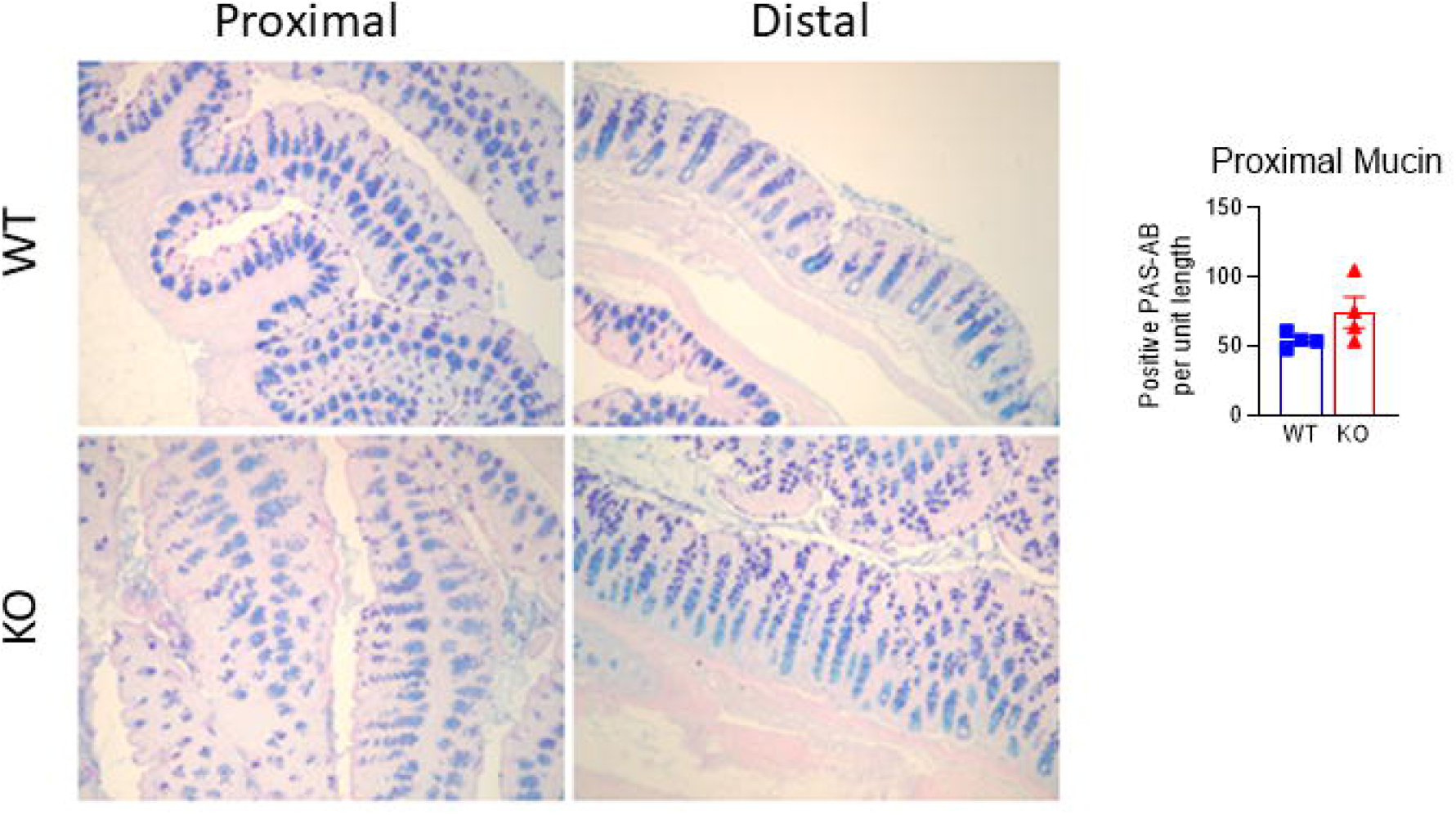
Mucin content is greater in the colons of PGDH KO versus WT mice. Representative images of PAS/Alcian Blue staining for proximal and distal mucin are shown. Quantification of proximal mucin content is graphed. Mean±SEM, N = 4 mice/group. **P=0.008 in distal colon by unpaired t test.

**Supplementary Figure 8.**
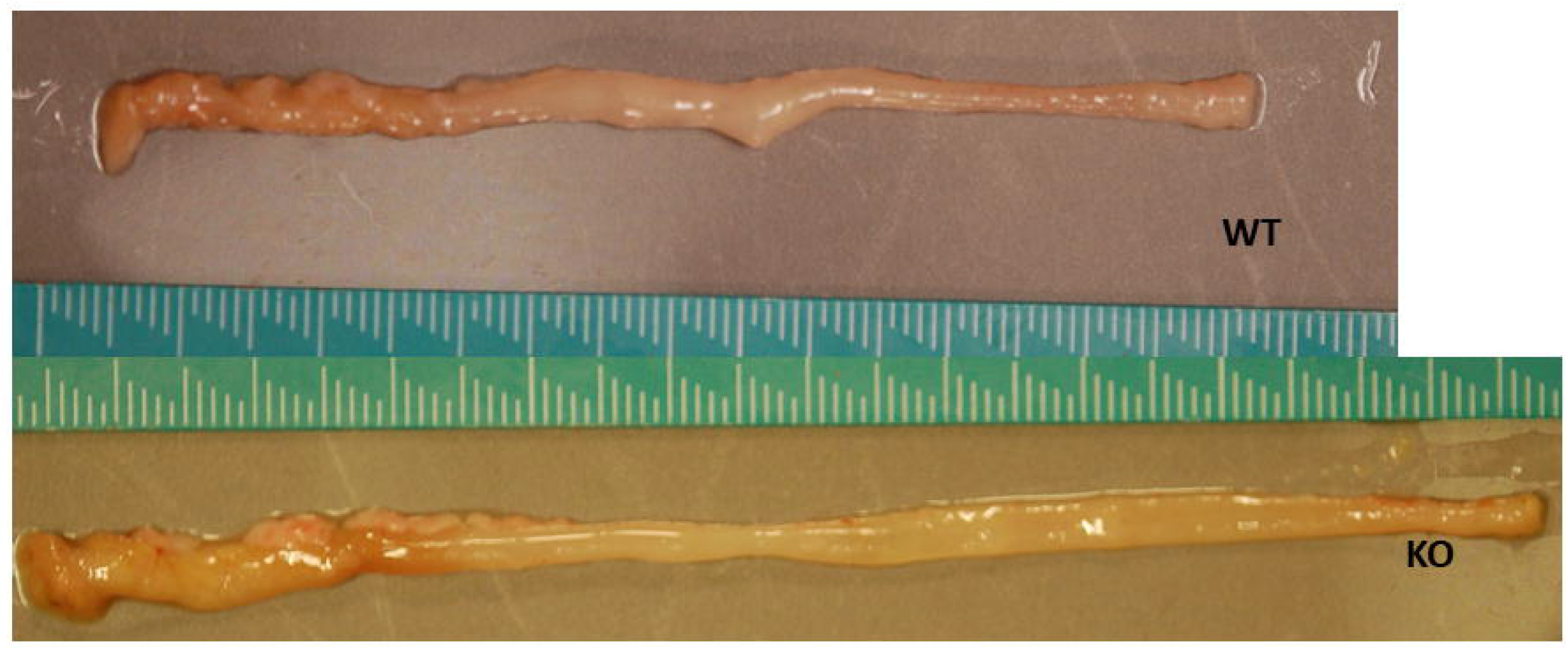
Gross colon morphology at day 10 of DSS colitis. A representative photograph of gross colons at day 10 of DSS colitis are shown for WT and PGDH KO.

**Supplementary Figure 9.**
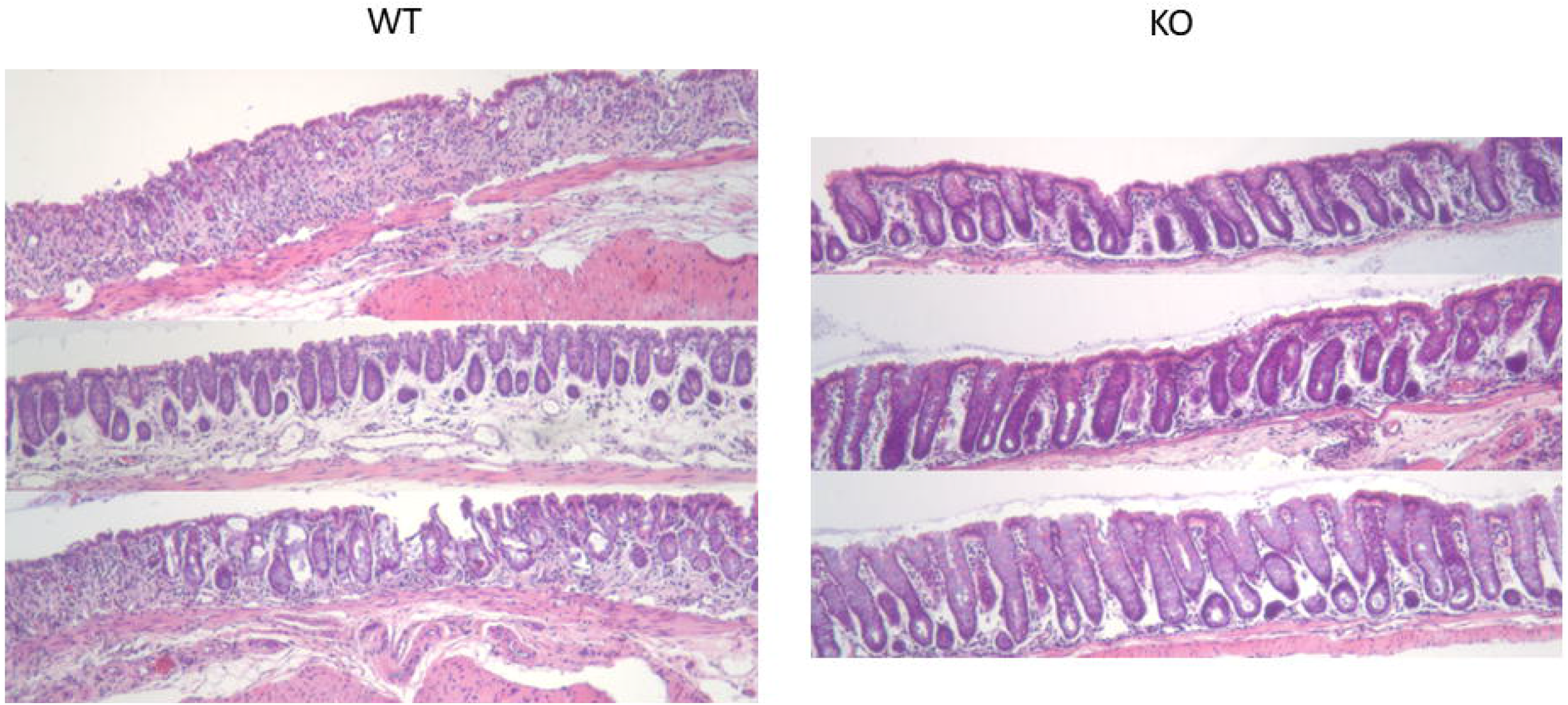
Distal colon histology at day 10 of DSS colitis. Additional representative distal colon cryptitis micrographs at day 10 of DSS colitis are shown for WT and PGDH KO.

**Supplementary Figure 10.**
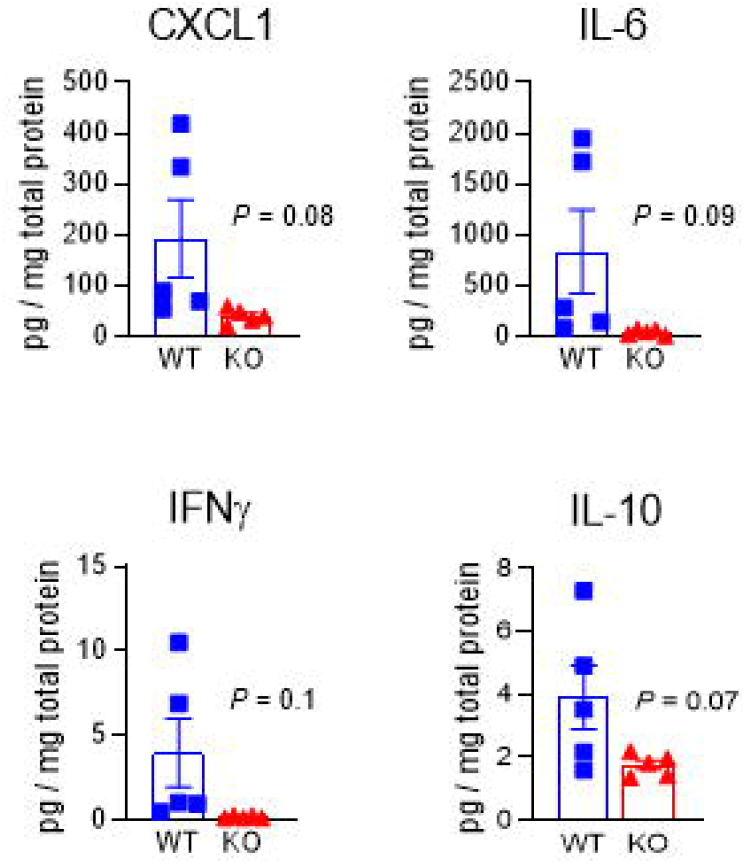
Inflammatory cytokine levels are attenuated in the colons of aged PGDH KO mice upon being injured by DSS. Quantification of CXCL1, IL6, IFN*γ*, and IL10 in the distal colons (pg/mg total protein) at day 10 of DSS colitis are compared. Mean±SEM, N = 5, P values obtained by unpaired t-tests.

